# Effect of Temperature on Gene Expression of *Escherichia marmotae*

**DOI:** 10.64898/2026.08.25.747177

**Authors:** Pelumi M. Oladipo, Ali M. Jomaa, Xiangmin Zhang, Jeffrey H. Withey, Jeffrey L. Ram

**Author notes:** corresponding author: Pelumi M. Oladipo, Jeffrey Ram.

## Abstract

Increased temperature is one of the first environmental cues encountered by bacteria upon entering a mammalian host. Here, we investigated the effects of two incubation temperatures on the transcriptome and proteome of *E. marmotae* and *E. coli*. As previous studies demonstrated that temperature affects motility in *E. marmotae*, our goal was to determine how temperature alters global gene expression at 37 °C versus 28 °C, and whether this response is conserved in *E. coli*. To do this, we grew strains of each species in static conditions at 28 °C and 37 °C and assessed gene expression by RNA transcriptome analysis and protein abundance by global proteomics.

Temperature altered the expression of 111 genes (2.7% of the genome analyzed) in *E. marmotae* and 99 genes (2.5%) in *E. coli* (adjusted p < 0.05, ≥2-fold change), with differential expression concentrated within specific functional pathways rather than reflecting global transcriptome-wide shifts. Comparable proportions of each proteome were similarly affected. In *E. marmotae*, genes spanning the class II and class III flagellar hierarchy and chemotaxis genes, along with operons for cellulose-dependent biofilm formation and nitrate respiration, were markedly downregulated at 37 °C. In contrast, genes associated with fimbrial adhesion and immune evasion, including *fimA/fimB*, *ompT*, and prophage-associated loci, were upregulated. Proteomic analysis corroborated these trends, showing coordinated loss of flagellar and chemotaxis proteins at 37 °C and increased abundance of stress-adaptation and host-interaction proteins, including OmpT. *E. coli* showed a partially overlapping but distinct response, with stronger enrichment of metabolic and amino-acid biosynthesis pathways and minimal changes in motility regulation. Together, our studies demonstrate that motility in *E. marmotae* is temperature-dependent and may represent a mechanism for immune evasion within the host.

**Importance:** *Escherichia marmotae* is an emerging member of the genus *Escherichia* that has increasingly been associated with human infections, yet the mechanisms that enable its transition from environmental reservoirs to the mammalian host remain poorly understood. This study provides the first transcriptomic and proteomic characterization of the temperature response of *E. marmotae* and demonstrates that growth at host temperature (37 °C) selectively alters pathways associated with bacterial lifestyle and virulence. Most notably, the coordinated repression of flagellar motility and chemotaxis, together with increased expression of factors associated with adhesion, stress adaptation, and host interaction, suggests that temperature serves as an environmental signal that promotes adaptation to the host. Comparison with *E. coli* further revealed that this response is not broadly conserved, highlighting distinct temperature-dependent regulatory strategies in *E. marmotae*. These findings expand our understanding of the biology of this understudied pathogen and suggest that reduced motility at mammalian body temperature may contribute to host adaptation and immune evasion.

**Graphical abstract:** 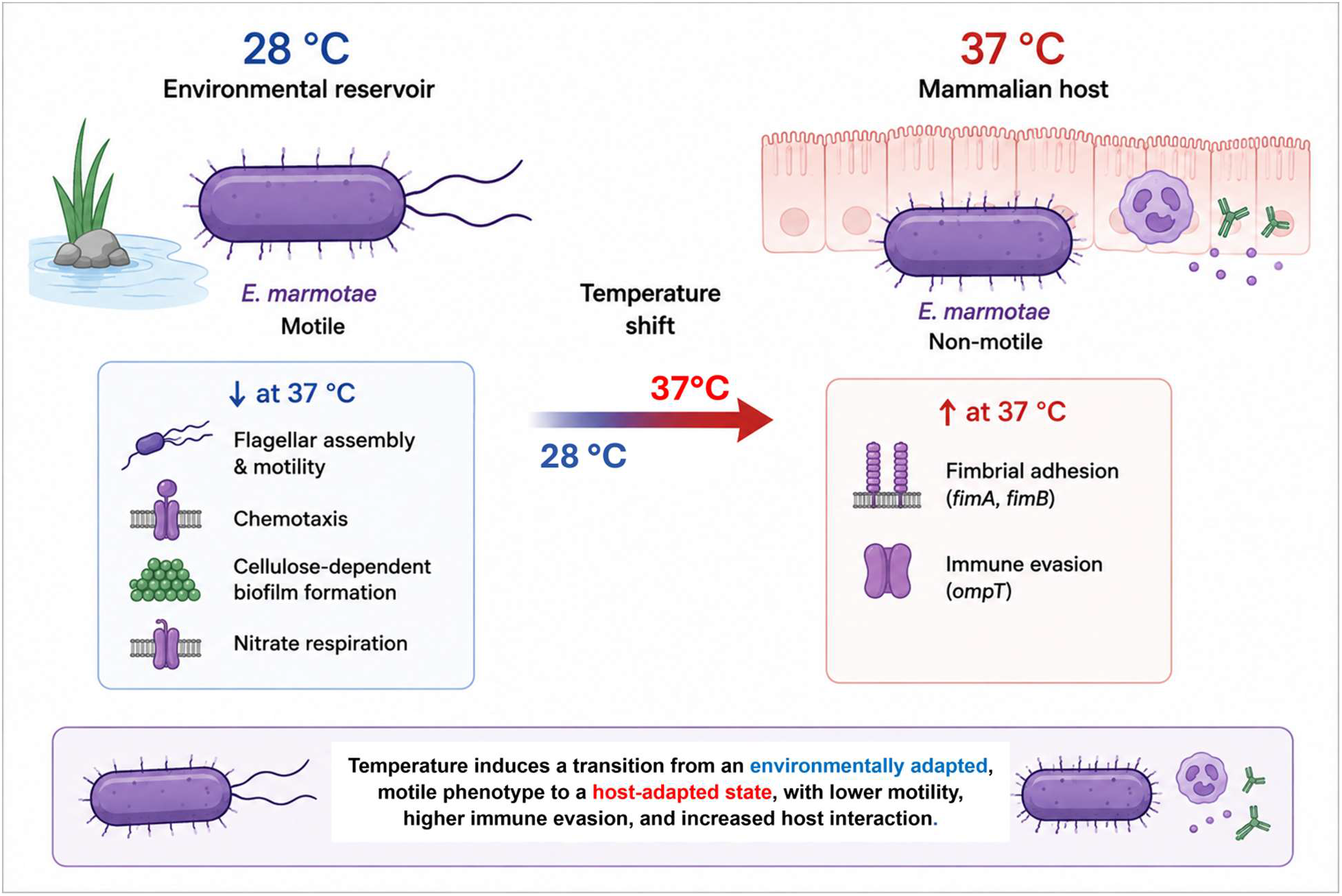

## 1.0 Introduction

The genus *Escherichia* comprises clinically and environmentally important members of the family *Enterobacteriaceae,* including *Escherichia coli* (*1*), *Escherichia fergusonii* (2), *Escherichia albertii* (3)*, Escherichia ruysiae* (4)*, Escherichia whittamii* (5), and *Escherichia marmotae* (6). *E. marmotae* was first identified as a cryptic *Escherichia* clade in 2009 (7) and subsequently as a distinct species in 2015 (6). Although *E. marmotae* was initially associated with animals and environmental reservoirs, accumulating reports now implicate *E. marmotae* in a range of human infections, including sepsis, urinary tract infections, spondylodiscitis, and pyelonephritis (8–11). These findings suggest that *E. marmotae* is not merely an environmental or commensal organism, but a bacterium with clinically relevant pathogenic potential. Nevertheless, despite growing evidence of its involvement in human disease, the molecular mechanisms underlying its pathogenicity remain poorly understood (12).

Recent genomic and phenotypic studies have identified putative virulence-associated determinants in *E. marmotae*. These include factors involved in motility gene regulation and structural proteins, type I fimbriae, enterobactins, heme uptake proteins, biofilm formation, iron siderophore and curli fimbriae (8, 9, 11, 13–17). Collectively, these features support the view that *E. marmotae* possesses a diverse repertoire of fitness and virulence traits that may facilitate host colonization, persistence, and infection. However, the expression and regulation of these determinants under different environmental conditions have been incompletely characterized.

Temperature is one of the most important environmental cues influencing bacterial physiology and virulence (18). For many pathogens, temperature shifts act as signals that indicate transition between environmental reservoirs and host-associated niches, thereby triggering coordinated changes in gene expression that optimize survival, colonization, and infection. While sudden temperature shifts often trigger protective stress responses, more subtle temperature differences can serve as signals that a bacterium has transitioned between environments or host niches (19). Pathogens may exploit these smaller temperature changes to coordinate the expression of key virulence determinants.

In *E. marmotae*, emerging evidence indicates that temperature plays an important regulatory role in motility. Previous work demonstrated significantly lower motility at 37 °C compared with 28 °C, as measured by decreased spread diameter in soft agar assays despite higher growth rate of *E. marmotae* at 37 °C (15). Consistent with this observation, reverse transcription quantitative PCR (RT-qPCR) analyses revealed reduced expression of key flagella-associated genes—*motA*, *fliC*, and *fliA*—at 37 °C relative to 28 °C (15). These findings suggest that temperature may play an important regulatory role in the expression of genes associated with motility and potentially other virulence-related pathways in *E. marmotae*.

However, current knowledge remains limited in several important ways. Previous investigations focused on only a small number of genes and did not address whether temperature-dependent regulation extends beyond flagellar function to broader cellular processes. In particular, the effects of temperature on other virulence-related pathways, including those involved in adhesion, biofilm formation, iron acquisition, stress adaptation, membrane functions, metabolism, and host interaction, are unknown. We hypothesized that other motility genes are downregulated at 37 °C in addition to those that were previously described (15) and also that temperature may cause differential expression of other functional categories of genes potentially involved in virulence.

This study is the first to investigate the effects of temperature on the expression of multiple genes in *E. marmotae* using high throughput transcriptomic and proteomic strategies and to compare *E. marmotae* and *E. coli* gene expression. The data described in this paper document the differential expression of numerous genes involved in motility and biofilm formation and other functions in *E. marmotae*, whose expression is affected by a change in incubation temperature between 28 °C and 37 °C. These findings enable a broader view of the expression of numerous motility-associated and other genes not previously investigated, in addition to supporting previous observations (15) of the changed expression of just a few motility-associated genes. Furthermore, by examining global changes in protein expression across these same temperature conditions, the functional implications of changes in gene transcription are further supported by changes measured in the abundance of specific proteins. This work aims to provide a broader understanding of the molecular pathways influenced by temperature and their possible contributions to the pathogenic potential of *E. marmotae*.

## 2.0 Materials and Methods

### 2. 1 Bacterial strains and growth conditions

Strains and plasmids used in this study are described in Table 1. All *E. marmotae* strains were cultured in Luria – Bertani (LB) medium. Individual colonies of three strains of *E. marmotae* and three strains of *E. coli* (Table 1) were picked, inoculated into 5 mL of LB broth, and incubated at 28 °C and 37 °C under static conditions for 18 h.

**Table 1:** Strains used in this study.

| Strain | Source |
| --- | --- |
| *E. marmotae* RAM 3032 | RamLab strains (15) |
| *E. marmotae* RAM 3054 | RamLab strains (15) |
| *E. marmotae* RAM 3024 | RamLab strains (15) |
| *E. coli* 430 | RamLab strains (20) |
| *E. coli* 250 | RamLab strains (20) |
| *E. coli* 218 | RamLab strains (20) |

### 2.2 RNA analysis

The overall RNA analysis workflow is illustrated in Figure 1. Details of growth conditions, RNA extraction methods, and subsequent high-throughput (RNA-seq) sequencing and bioinformatics analysis are given below.

**Figure 1:**
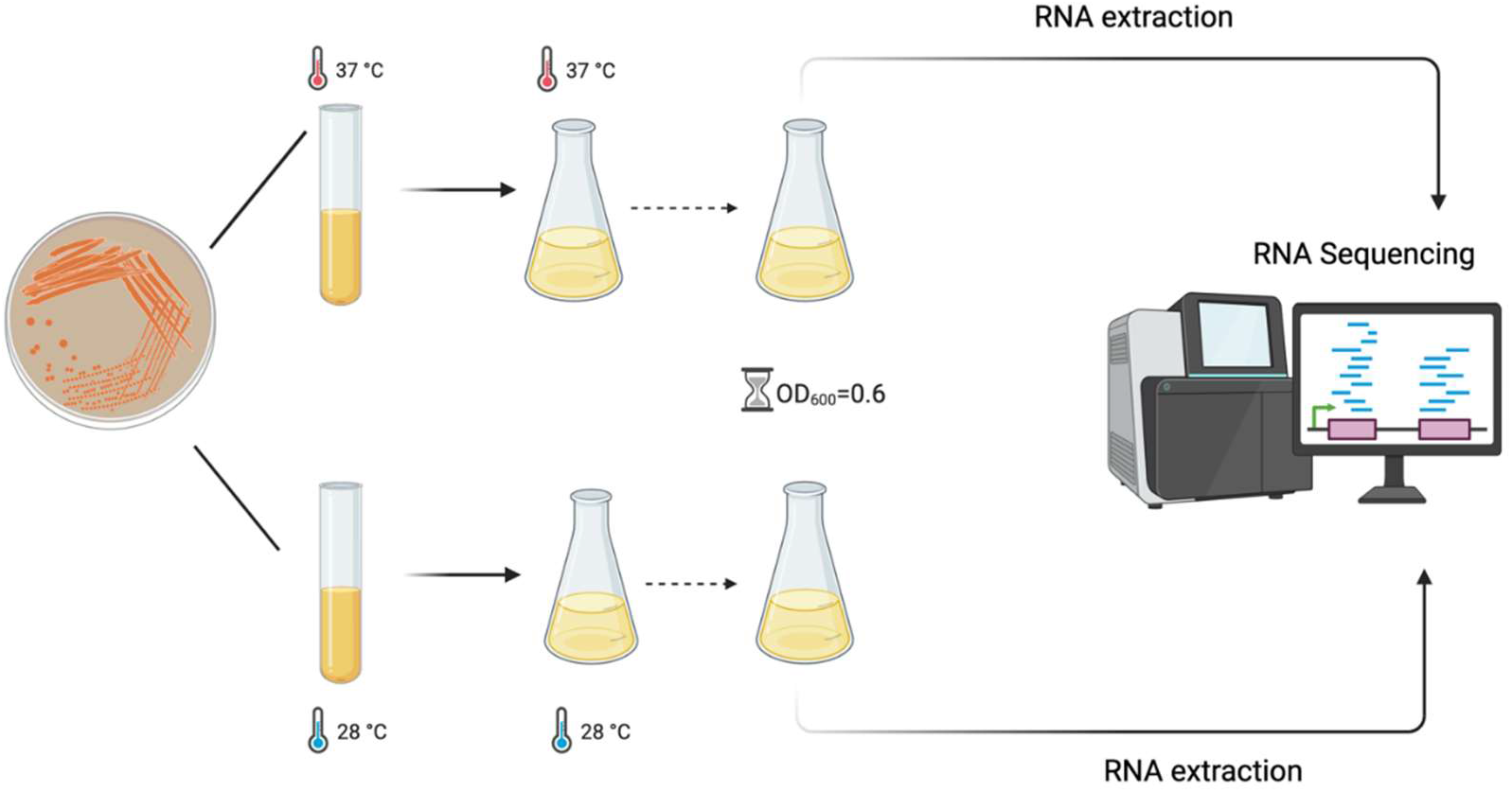
Diagram of the experimental setup for RNA-seq of *E. marmotae* and *E.coli* grown at two temperatures. Bacterial cultures were grown at 28 °C and 37 °C to mid-log phase (OD_600_ = 0.5 - 0.6). Total RNA was extracted and prepared for high-throughput sequencing to compare temperature–dependent gene expression profiles.

#### 2.2.1. RNA extraction

After culture at 28 °C and 37 °C under static conditions for 18 h incubation, the overnight cultures were diluted 1:100 in fresh LB broth, incubated further, and harvested at an optical density (OD_600_) of 0.5 - 0.6. The bacteria were pelleted by centrifugation at 5,000 X g for 10 min, and RNA was extracted with an RNeasy kit (Qiagen, Germany) using a modified protocol (21). The modifications included adding 700 µL of Buffer RW1 to the RNeasy spin column before adding the DNase I stock solution. All centrifugation steps were performed for 30 seconds at >8000 x g (12,000 rpm). Purified RNA was collected by placing the spin column in a new collection tube and centrifuging for 1 minute at >8000 x g (14,500 rpm). The integrity of the purified RNA was examined by agarose gel electrophoresis, visualizing the 23S/16S banding pattern. RNA concentration was measured using a NanoDrop 2000 spectrophotometer (Thermo Scientific, USA).

The use of three strains of *E. marmotae* (RAM 3032, RAM 3024, and RAM 3054, isolated as described by (15)), and three strains of environmental *E. coli* (430, 218, and 250, isolated as described by (20)) in this study served as representative independently isolated biological replicates of each species. In addition, two independently initiated cultures of each strain grown under each growth condition (28 °C and 37 °C) were analyzed to ascertain reproducibility of the observed gene expression patterns.

#### 2.2.2. RNA Sequencing, Read Processing, and Differential Expression Analysis

Library construction and RNA sequencing were performed on the Illumina platform (2 × 150 bp paired-end reads) by Azenta Life Sciences (Burlington, MA and South Plainfield, NJ, USA). FASTQ sequence files were processed, and expression profiles were represented as read counts. Adapter sequences and low-quality bases were trimmed using cutadapt v 5.2 (22) and Trimmomatic v0.36, retaining reads with a paired quality score >30 across all runs. High-quality reads were aligned using Bowtie2 v2.5.5 to the *Escherichia marmotae* RAM 3032 reference genome (GenBank accession: <u>JBNVMU000000000</u>) using standard alignment workflows. For *E. coli,* high quality reads were aligned to the whole genome sequence of *E. coli* EC96 (GenBank accession: CP060748.1).

Read counts were quantified with featureCounts (Subread v1.5.2), and expression levels were normalized as fragments per kilobase of transcript per million mapped reads (FPKM), taking into account both transcript length and sequencing depth. Unique gene hit counts were used as the input for differential expression analysis.

Differential gene expression analysis was conducted using the DESeq2 package v1.50.2 (23). A Wald test was applied to estimate p-values and log₂ fold changes between defined experimental groups. Differential expression was considered to be significant for genes whose transcripts exhibited an absolute log₂ fold change > 1 (i.e., at least doubling of expression) and an adjusted p-value < 0.05. MA plots (Bioconductor, Vienna, Austria) were used to visualize differential effects of temperature on gene expression across a range of gene abundances.

For testing multiple genes simultaneously, the *p*-values were further corrected using the Benjamini-Hochberg false discovery rate (FDR) method (24). Venn diagrams were generated to compare transcribed genes across different conditions. For principal component analyses (PCA), mapped reads were variance stabilizing transformation (VST) normalized using the *vst* function of DESeq2 and subsequently used by the *plotPCA* function of DESeq2 to produce the PCA, which was further visualized using ggplot2.

### 2.3. Protein analysis

#### 2.3.1. Protein Extraction

After culture of the same strains as were used for RNA analysis at 28 °C and 37 °C under static conditions for 18 h, lysates were prepared from intact bacterial cells from 30 mL of bacterial culture. The culture was centrifuged at 5,000 × *g* for 15 min, and the cell pellet was washed twice with PBS. Bacterial cell pellets were resuspended in 400 µL of 2.5% LiDS, then were heated to 95 °C for 5 minutes then filtered through spin columns (Pierce #89868) to produce a clear lysate solution. An aliquot of each filtered lysate was taken for protein analysis and the remainder of each sample was buffered with 20 mM Tris pH 8.5. The remaining material was reduced with 5 mM DL-dithiothreitol (DTT, Sigma, cat. no. D5545) for 30 mins at 37 °C, alkylated with 15 mM iodoacetamide (IAA, Sigma, cat. no. I1149) for 30 mins at room temperature in the dark, and quenched by the addition of 5 mM DTT. For each sample, 20 µg of protein was transferred to a new tube and adjusted to a final volume of 50 µL. Samples were acidified by addition of 5 µL of 12% phosphoric acid then proteins precipitated by addition of 350 µL of 90% MeOH (Methanol) in 100 mM Triethylammonium bicarbonate (TEAB). Protein pellets were washed with 500 µL of 80% MeOH in 10 mM TEAB, air-dried, and resuspended in 30 µL of 20 mM Tris (pH 8.0), 10 mM CaCl_2_, and 10% acetonitrile (ACN). Trypsin (Promega, cat. no. V5113) was added at 1 µg per sample, and digestion was carried out at 47 °C for 2 hr. Samples were dried and reconstituted in 0.1% formic acid (FA).

#### 2.3.2. Mass spectrometry analysis

LC-MS/MS analysis utilized a Thermo Scientific Vanquish-Neo chromatography system with an Acclaim PepMap 100 trap column (100 µm × 2 cm, C18, 5 µm particle size, 100Å pore size), and an Ion Optics Aurora column (75 µm × 25 cm) maintained at 45 °C in an Easy Spray source. A 120-minute gradient was applied, starting with 1% of solution of B (80% ACN, 0.1% FA), and increasing to a final 42% of solution B. Data-independent analysis (DIA) was performed on a Thermo Scientific Orbitrap Eclipse mass spectrometer. MS1 spectra are acquired at 120,000 resolution over a mass range of 350–1,000 *m/z*, with an automatic gain control (AGC) target of 3 × 10⁶. MS2 spectra were acquired in the Orbitrap at 15,000 resolution, with a 200-1600 Da window. Fragmentation for MS2 spectra was completed using 15 Da windows between 350 to 900 Th with HCD fragmentation at a collision energy of 32, a maximum injection time of 80 msec, and an AGC target of 3e6.

#### 2.3.3. Protein identification and quantification

Mass spectrometry data were processed using Spectronaut version 19.3 (Biognosys) with factory default settings. Protein identification was performed against either the *E. coli* or *E. marmotae* UniProt database, as appropriate. Trypsin was specified as the proteolytic enzyme, allowing up to two missed cleavages. Variable modifications included methionine oxidation and protein N-terminal modifications, including acetylation, methionine loss, and methionine loss with acetylation. Carbamidomethylation of cysteine residues was specified as a fixed modification. For the entire dataset, false discovery rate (FDR) thresholds were set at 1% for precursor identification and 1% for peptide identification. Proteomic data were subsequently analyzed for potential altered gene groups by comparison to gene ontology groups. The mass spectrometry proteomics data have been deposited to the ProteomeXchange Consortium via the PRIDE repository under accession number PXD081413 (25).

#### 2.3.4. Functional Enrichment and Network Analysis

Gene Set Enrichment Analysis (GSEA) was conducted using the clusterProfiler package V4.18.4 to determine significantly enriched COG (Clusters of Orthologous Groups) functions and categories (26). Differentially expressed genes with valid gene symbols were mapped to Entrez Gene IDs using the org.EcK12.eg.db annotation database v3.22.0. Gene Ontology (GO) enrichment was assessed across Biological Process, Molecular Function, and Cellular Component categories, with adjusted p-values used to rank enriched terms.

Protein–Protein Interaction (PPI) networks were explored using the STRING database v12.0 (27) and accessed at https://string-db.org/. Components of these PPI networks of significantly regulated genes were mapped to STRING identifiers for *E. coli* K-12 reference species (taxonomy ID: 511145), using a medium-confidence interaction score threshold of 400.

## 3.0 Results

### 3.1 RNA-seq analysis to identify temperature -regulated genes in *E. marmotae*

Cultures of *E. marmotae* and *E. coli* isolates at 28 °C and at 37 °C, followed by RNA extraction and RNA transcript analysis on the Illumina platform, resulted in a large number of quality reads for all extracts, enabling the analysis of differential gene expression between the two culture temperatures in each of the *Escherichia* species. After adapter trimming and quality filtering, the total numbers of reads, averaged across the biological replicates for each condition were as follows: *E. marmotae* grown at 28 °C, 38,013,248 reads; *E. marmotae* grown at 37 °C, 38,180,782 reads; *E. coli* grown at 28 °C, 38,203,886 reads; and *E. coli* grown at 37 °C, 31,834,381 reads. This process resulted in the identification of transcripts of 4131 genes for *E. marmotae* and 3964 genes for *E.coli* suitable for statistically reliable differential gene analysis. Detailed per-sample read quality, trimming, and alignment metrics are given in Table 2.

**Table 2:**
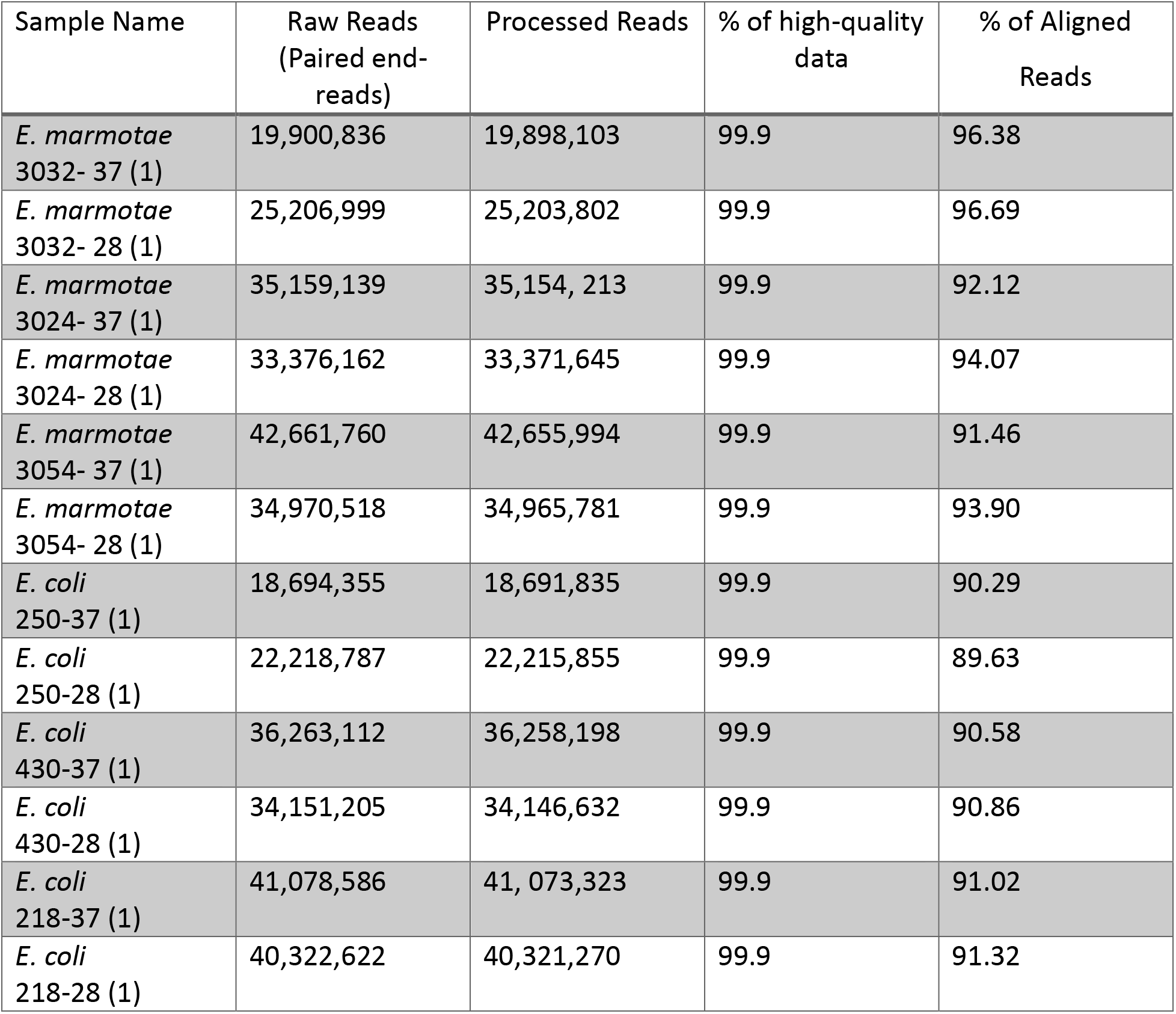
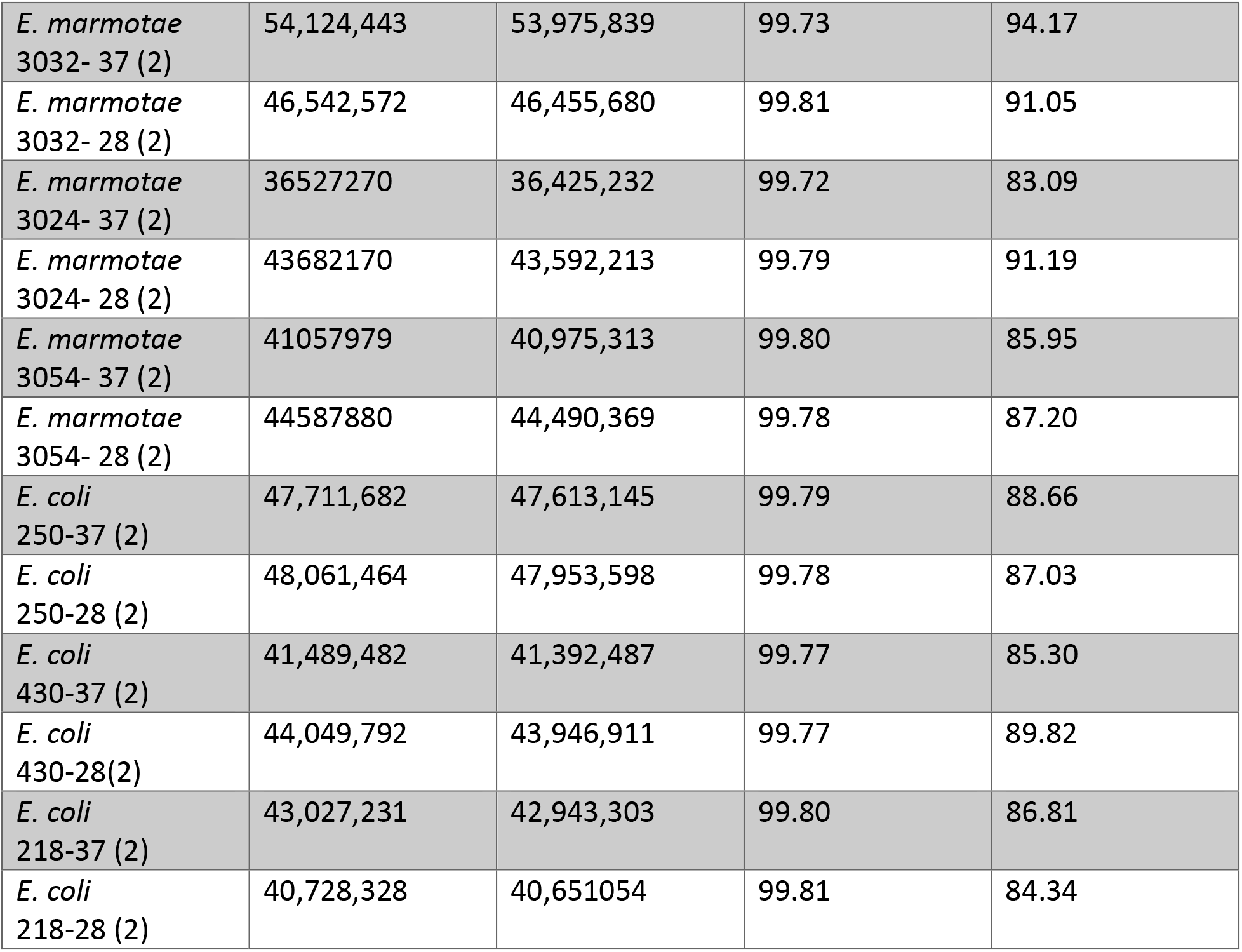
Overview of raw reads and mapped sequences for all the samples. Three strains of each species were analyzed, with two independently initiated replicate cultures per strain under each temperature condition. Numbers in parentheses following the sample names indicate the replicate culture number, where (1) denotes the first culture replicate and (2) denotes the second culture replicate for that strain.

#### 3.1.1. Differential Expression analysis

Differentially expressed genes were defined as those with an adjusted p-value < 0.05 and an absolute log₂ fold change ≥ 1. Genes with log₂ fold change > 1 were classified as upregulated at 37 °C, while genes with log₂ fold change < −1 were classified as downregulated at 37 °C. Genes with log₂ fold change values between −1 and 1 were considered not strongly differentially regulated.

In *E. marmotae*, temperature affected the expression of 111 genes by at least two-fold, representing approximately 2.7% of the genes included in the analysis (Figure 2; Table 3, Table S1). Of these, 47 genes were upregulated, and 64 genes were downregulated at 37 °C relative to 28 °C. In *E. coli*, 99 genes were differentially expressed, representing approximately 2.5% of the genes included in the analysis (Figure 3, Table 4, Table S2). Among these, 42 genes were upregulated, and 57 genes were downregulated at 37 °C relative to 28 °C.

**Figure 2:**
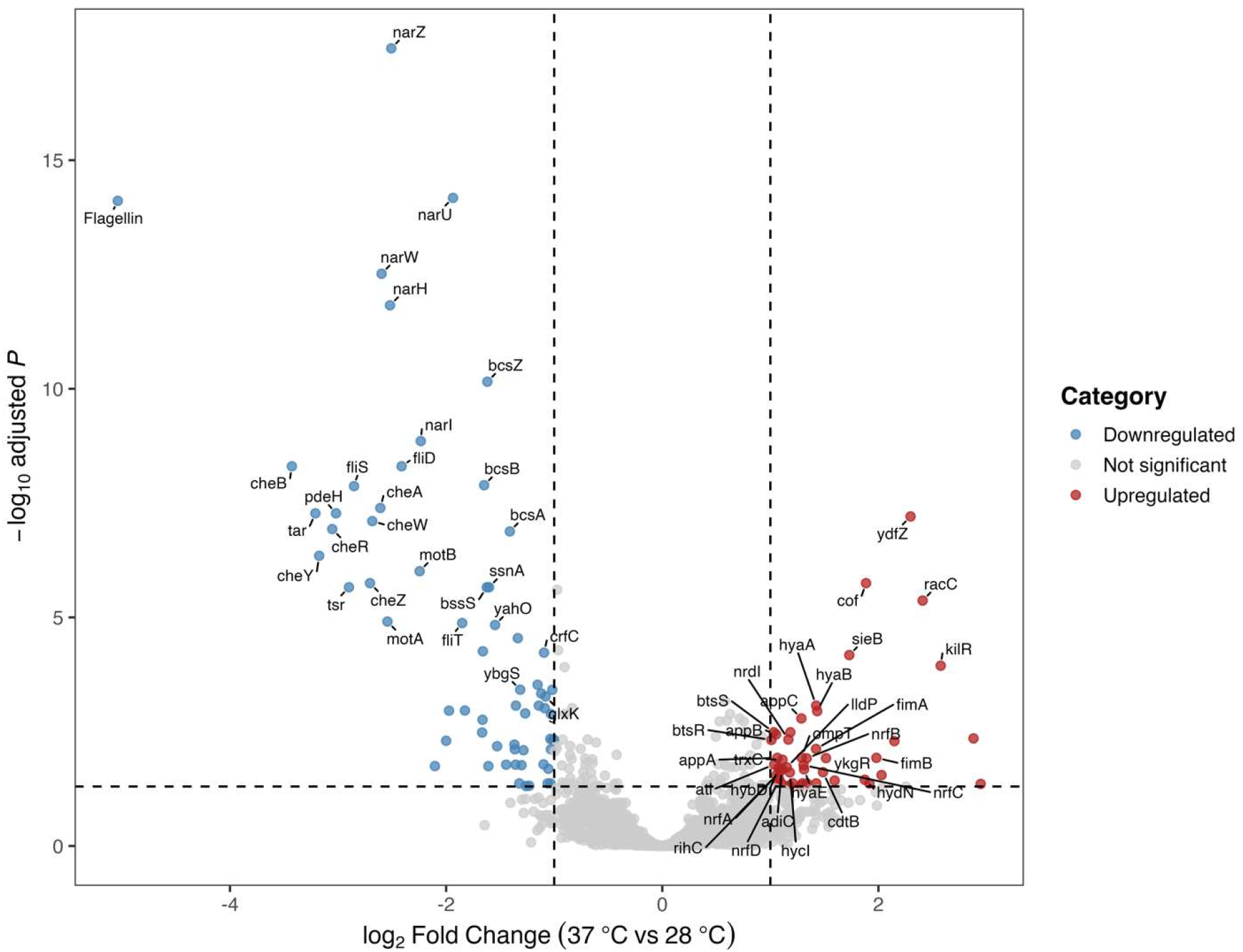
Volcano plot of differential gene expression in *E. marmotae* at 37 °C relative to 28 °C. Each point represents a gene with detectable expression. Red and blue points indicate genes significantly upregulated and downregulated, respectively, at 37 °C, based on an adjusted p-value < 0.05 and an absolute log₂ fold change ≥ 1. Grey points represent genes that were not significantly differentially expressed.

**Figure 3:**
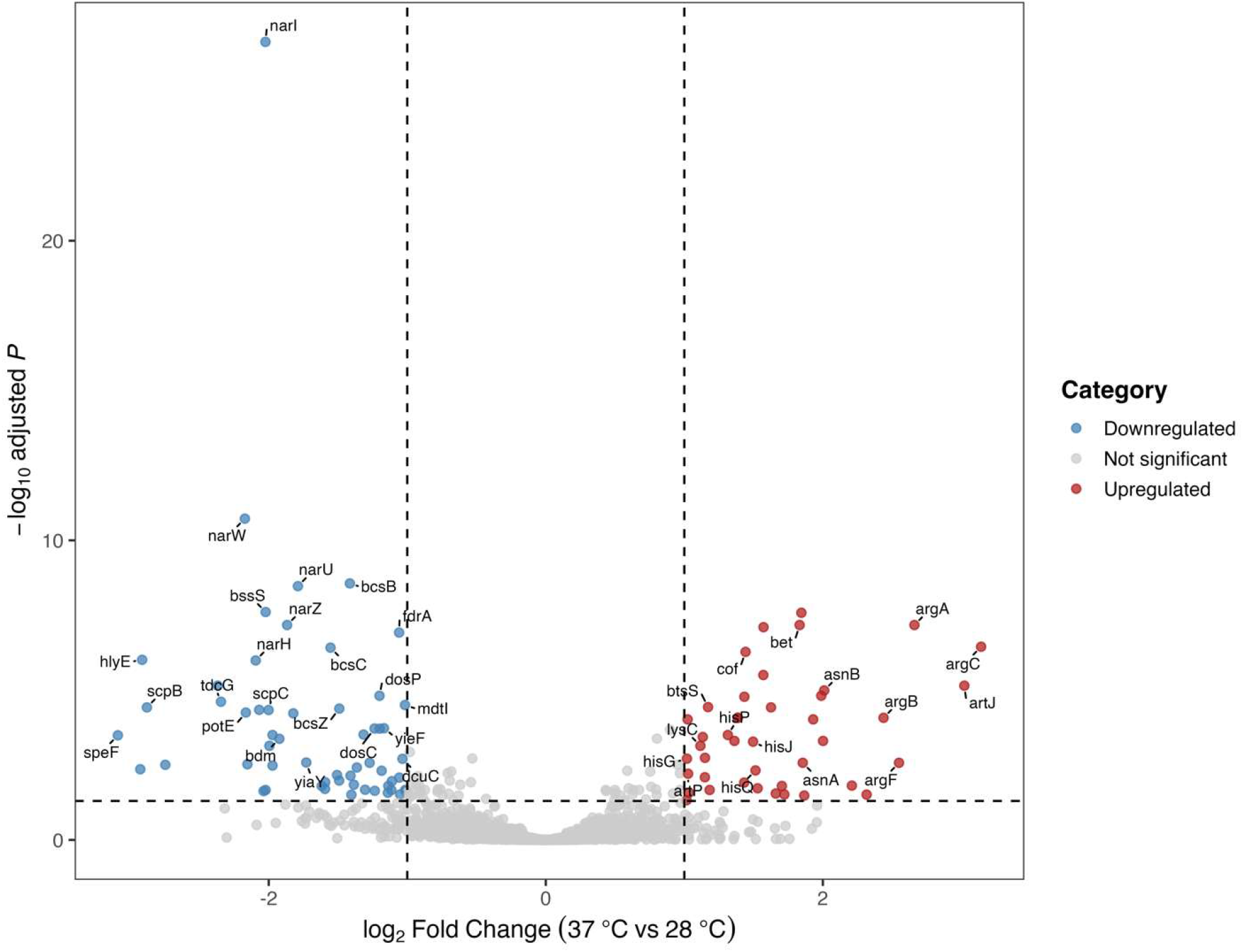
Volcano plot of differential gene expression in *E. coli* at 37 °C relative to 28 °C. Each point represents a gene with detectable expression. Red and blue points indicate genes significantly upregulated and downregulated, respectively, at 37 °C, based on an adjusted p-value < 0.05 and an absolute log₂ fold change ≥ 1. Grey points represent genes that were not significantly differentially expressed.

**Table 3:**
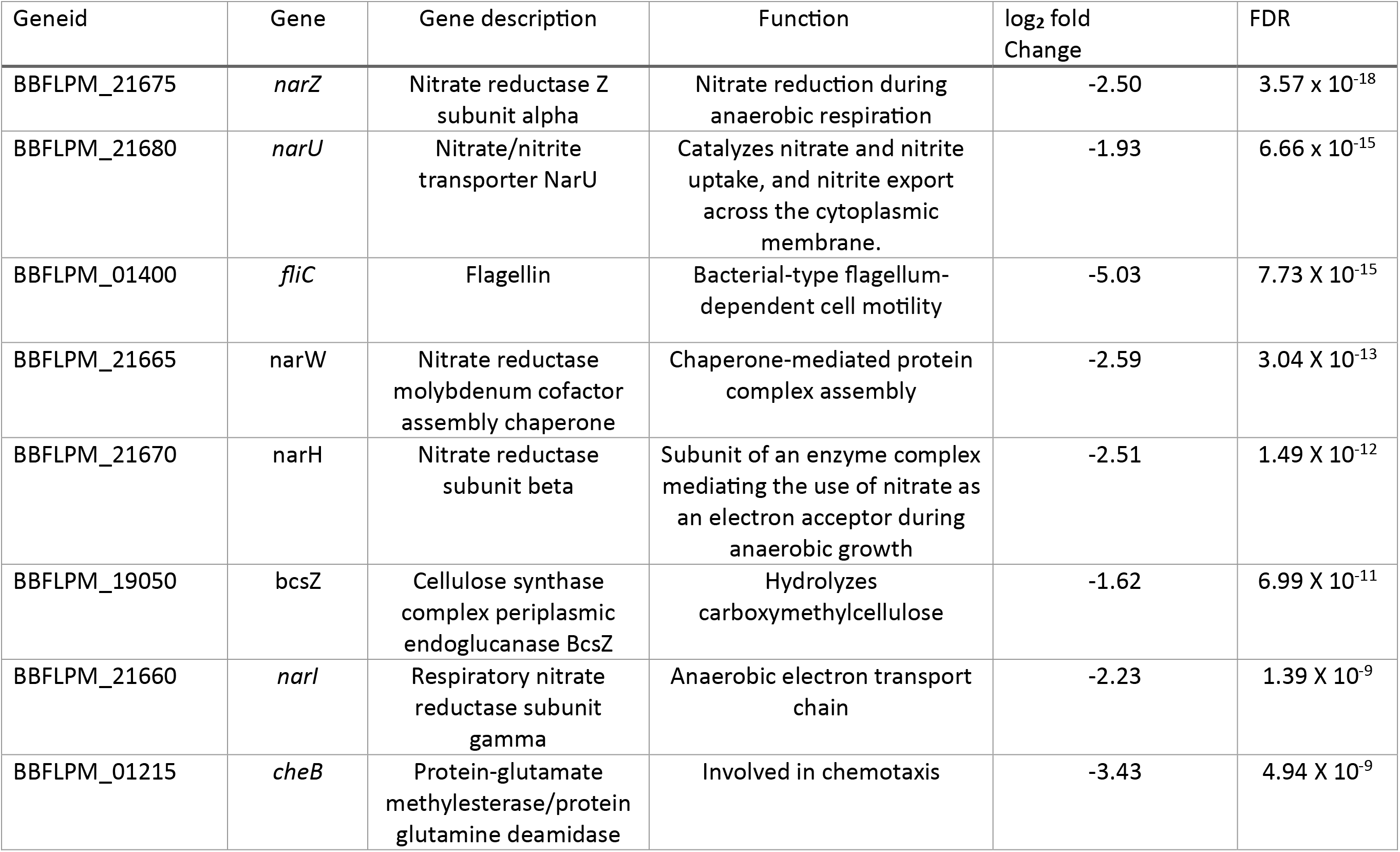
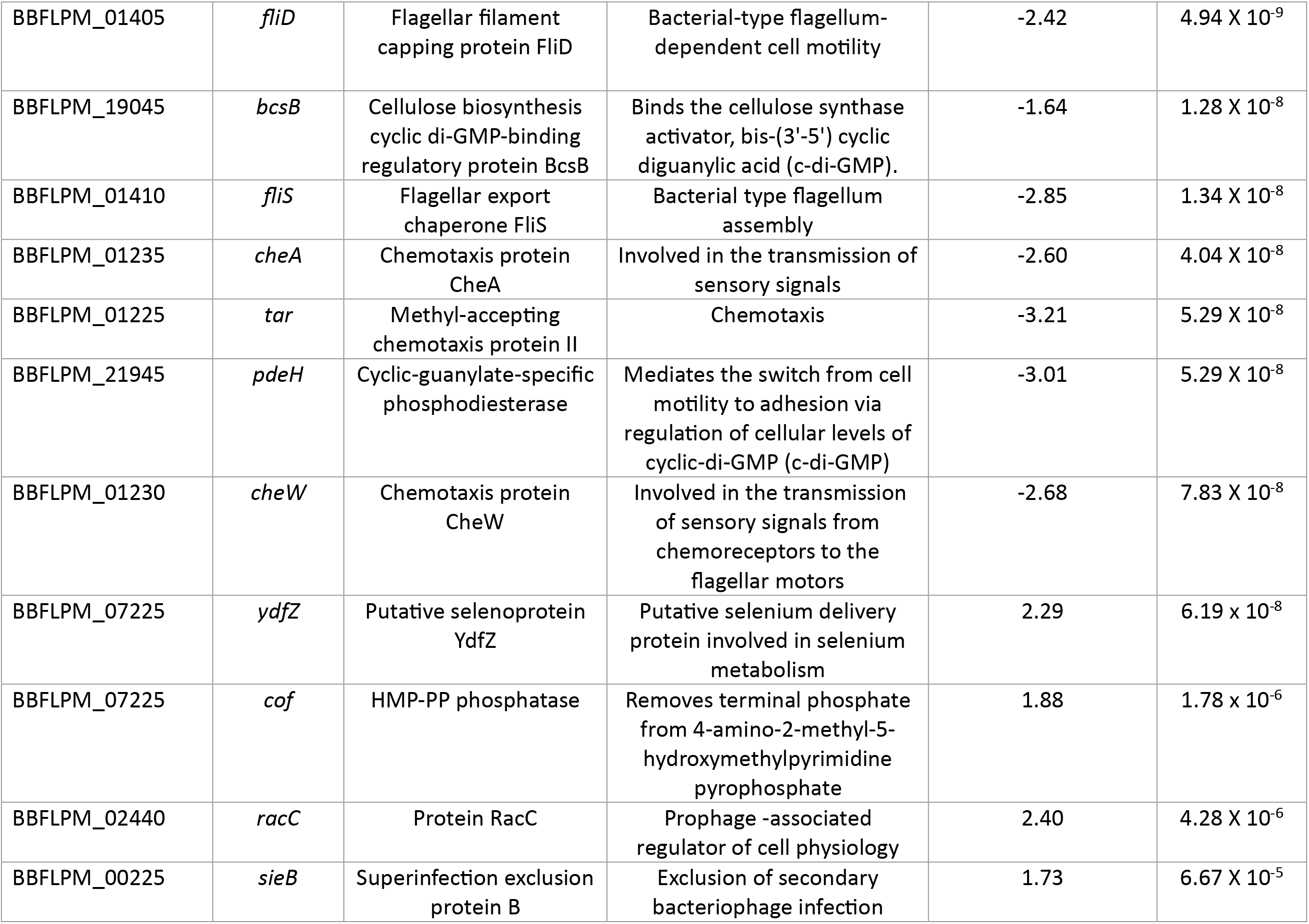
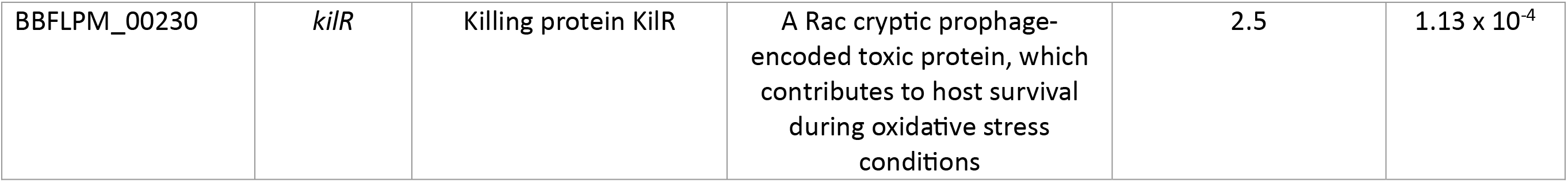
Top 20 differentially expressed genes in *E. marmotae* at 37°C relative to 28°C. Positive log₂ fold change values indicate higher expression at 37°C, whereas negative values indicate lower expression at 37°C. FDR = Benjamini–Hochberg false discovery rate

**Table 4:** Top 20 differentially expressed genes in *E. coli* at 37 °C relative to 28 °C. Positive log₂ fold change values indicate higher expression at 37°C, whereas negative values indicate lower expression at 37°C. FDR = Benjamini–Hochberg false discovery rate.

| Geneid | Gene | Gene description | Function | log<sub>2</sub> fold change | FDR |
| --- | --- | --- | --- | --- | --- |
| IAI02_11555 | *narI* | Respiratory nitrate reductase subunit gamma | The nitrate reductase enzyme complex allows *E.coli* to use nitrate as an electron acceptor during anaerobic growth. | -2.02 | 2.31 x 10<sup>-27</sup> |
| IAI02_11550 | *narW* | Nitrate reductase molybdenum cofactor assembly chaperone | Chaperone-mediated protein complex assembly | -2.17 | 1.88 x 10<sup>-11</sup> |
| IAI02_00630 | *bcsB* | Cellulose biosynthesis cyclic di-GMP-binding regulatory protein BcsB | Binds the cellulose synthase activator, bis-(3'-5') cyclic diguanylic acid (c-di-GMP). | -1.41 | 2.74 x 10<sup>-9</sup> |
| IAI02_11535 | *narU* | Nitrate/nitrite transporter | Catalyzes nitrate and nitrite uptake and nitrite export across the cytoplasmic membrane. | -1.78 | 3.38 x 10<sup>-9</sup> |
| IAI02_13470 | *BssS* | Biofilm formation regulator BssS | Regulation of single -species biofilm formation | -2.02 | 2.47 x 10<sup>-8</sup> |
| IAI02_05055 | *argA* | Amino-acid N-acetyltransferase | L-arginine biosynthetic process | 2.66 | 6.70 x 10<sup>-8</sup> |
| IAI02_13050 | QNN15195.1 | Hypothetical protein | Hypothetical protein | 1.84 | 2.61 x 10<sup>-8</sup> |
| IAI02_11540 | *narZ* | Nitrate reductase Z subunit alpha | Nitrate reduction during anaerobic respiration | -1.86 | 6.70 x 10<sup>-8</sup> |
| IAI02_16125 | *Bet* | Phage recombination protein Bet | DNA recombination | 1.83 | 6.70 x 10<sup>-8</sup> |
| IAI02_16160 | QNN15755.1 | DUF4222 domain-containing protein | DUF4222 domain-containing protein | 1.57 | 7.92 x 10<sup>-8</sup> |
| IAI02_17250 | *fdrA* | Acyl-CoA synthetase FdrA | Fatty acid metabolism | -1.05 | 1.19 x 10<sup>-7</sup> |
| IAI02_20120 | *argC* | N-acetyl-gamma-glutamyl-phosphate reductase | Arginine biosynthesis | 3.14 | $3.55 \times 10^{-7}$ |
| IAI02_00640 | *bcsC* | Cellulose biosynthesis protein BcsC | Required for maximal bacterial cellulose synthesis | -1.55 | $3.82 \times 10^{-7}$ |
| IAI02_16635 | *cof* | HMP-PP phosphatase | Antibiotic catabolic process | 1.44 | $5.28 \times 10^{-7}$ |
| IAI02_12705 | *hlyE* | Hemolysin HlyE | Toxin that has some hemolytic activity towards mammalian cells | -2.91 | $9.71 \times 10^{-7}$ |
| IAI02_11545 | *narH* | Nitrate reductase subunit beta | The nitrate reductase enzyme complex allows *E.coli* to use nitrate as an electron acceptor during anaerobic growth | -2.09 | $1.01 \times 10^{-6}$ |
| IAI02_03470 | QNN17323.1 | Enamine/imine deaminase | Amino acid metabolism | -2.36 | $7.07 \times 10^{-6}$ |
| IAI02_14485 | *artJ* | ABC transporter substrate-binding protein ArtJ | L-arginine import across plasma membrane | 3.02 | $7.10 \times 10^{-6}$ |
| IAI02_15425 | *asnB* | Asparagine synthase B | Catalyzes the ATP-dependent conversion of aspartate into asparagine, using glutamine as a source of nitrogen | 2.00 | $1.02 \times 10^{-5}$ |
| IAI02_11430 | *dosP* | oxygen-sensing cyclic-di-GMP phosphodiesterase | Heme-based oxygen sensor protein displaying phosphodiesterase (PDE) activity toward c-di-GMP in response to oxygen availability | -1.20 | $1.53 \times 10^{-5}$ |

Differentially expressed genes were distributed across a broad range of transcript abundances in both species. Expression differences at the two temperatures is represented by the MA plot in Figure 4, which graphs differences in the log_2_ transcript counts between 37 °C and 28 °C versus the mean abundance of the transcripts. This analysis shows that temperature-dependent regulation was not restricted to highly or weakly expressed genes but occurs across the entire range of abundances for both species.

**Figure 4:**
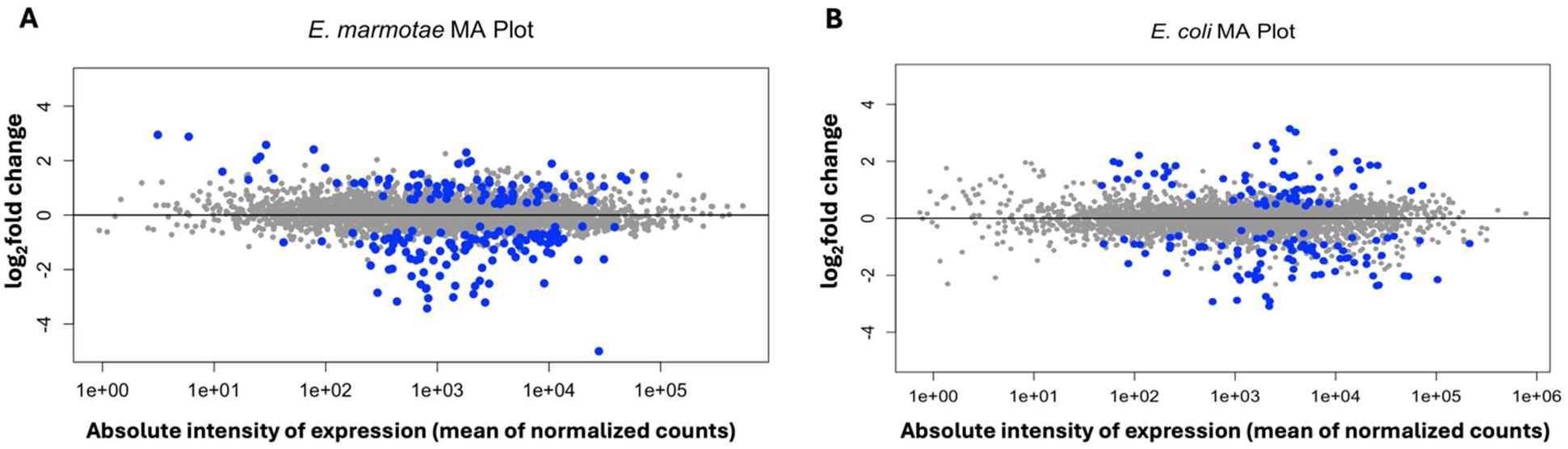
Differential effect of temperature on transcript abundance versus mean transcript abundance in (A) *E. marmotae* and (B) *E. coli*, displayed as MA plots. Each point represents the log₂ fold change in expression (vertical scale) of a gene at 37 °C relative to 28 °C plotted against the mean normalized abundance of that gene (horizontal axis). Blue points indicate significantly differentially expressed genes (adjusted p-value of <0.05), whereas grey points represent genes that were not significantly differentially expressed.

#### 3.1.2. Functional Roles of Thermo-Responsive Genes

Although distributed across a broad range of abundances, the temperature-elicited changes in gene expression nevertheless appear to focus on different genes in *E. marmotae* and *E. coli*. These differences in which genes are upregulated or downregulated between the two species are succinctly illustrated in the Venn diagram in Figure 5. Here, we see that *E. marmotae* and *E. coli* share only 9 out of 110 genes that are significantly downregulated when the culture temperature increases from 28 °C to 37 °C, and they share only 3 out of 84 genes that are significantly upregulated at the higher temperature. Moreover, no gene that was significantly downregulated in *E. marmotae* was significantly upregulated in *E. coli*, nor vice versa.

**Figure 5.**
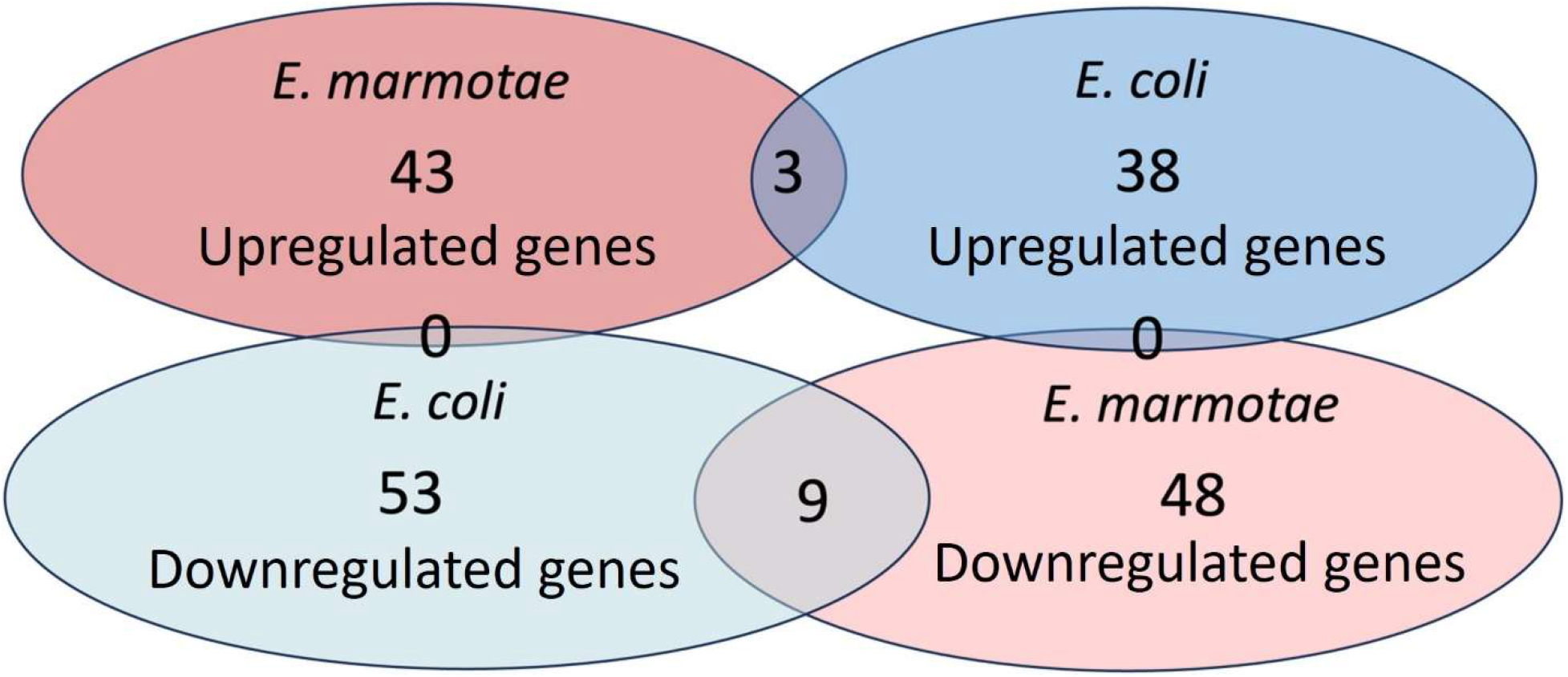
Venn diagram showing the overlap of significantly upregulated and downregulated genes in *E. marmotae* and *E. coli* at 37 °C relative to 28 °C. The diagram highlights genes uniquely or commonly regulated between species and direction of expression change, based on genes in Figures 2 and 3 with absolute log₂ fold changes ≥ 2 and adjusted p-values < 0.05.

Given the small amount of overlap between *E. marmotae* and *E. coli* in which genes are upregulated or downregulated, we analyzed whether *E. marmotae* has unique functional groups of genes that are affected by temperature, contrasting with those affected by temperature in *E. coli*. An initial analysis of the functions of the gene products of significantly changed transcripts in the volcano plot for *E. marmotae* (Figure 2) reveals that many of the downregulated genes in *E. marmotae* (20 genes out of 64 downregulated genes) are associated with motility and chemotaxis (*fliC, cheA/B/R/W/Y/Z, tar, tsr, motA/B, fliA/D/S/T*, and *flgK/L/M/N)* (15, 28). Additional downregulated genes include those involved in biofilm formation, such as *bcsA/B/Z*, *csgD/E/F/G*, and *bssS* (29), as well as genes linked to nitrate metabolism (*narZ/U/W/H/I*) (30). Other downregulated genes are involved in transport, envelope structure, and stress-response functions. Notably, flagellin showed the strongest decrease in expression, with an approximately 32-fold reduction at 37 °C relative to 28 °C. Flagellin was among the most statistically significant differentially expressed genes (FDR, 7.73 X 10^−15^; see Figure 2 and Table 3). Regarding temperature dependent upregulation, most genes whose expression increased at 37 °C were associated with adhesion and colonization, such *as fimA, fimB,* and *ompT*; expression of prophage or mobile element-associated genes such as *racC, sieB*, and *cdtB* also increased. These functional groupings of temperature-regulated genes in *E. marmotae* are further emphasized by formal Gene Ontology analysis (Figure 6).

**Figure 6.**
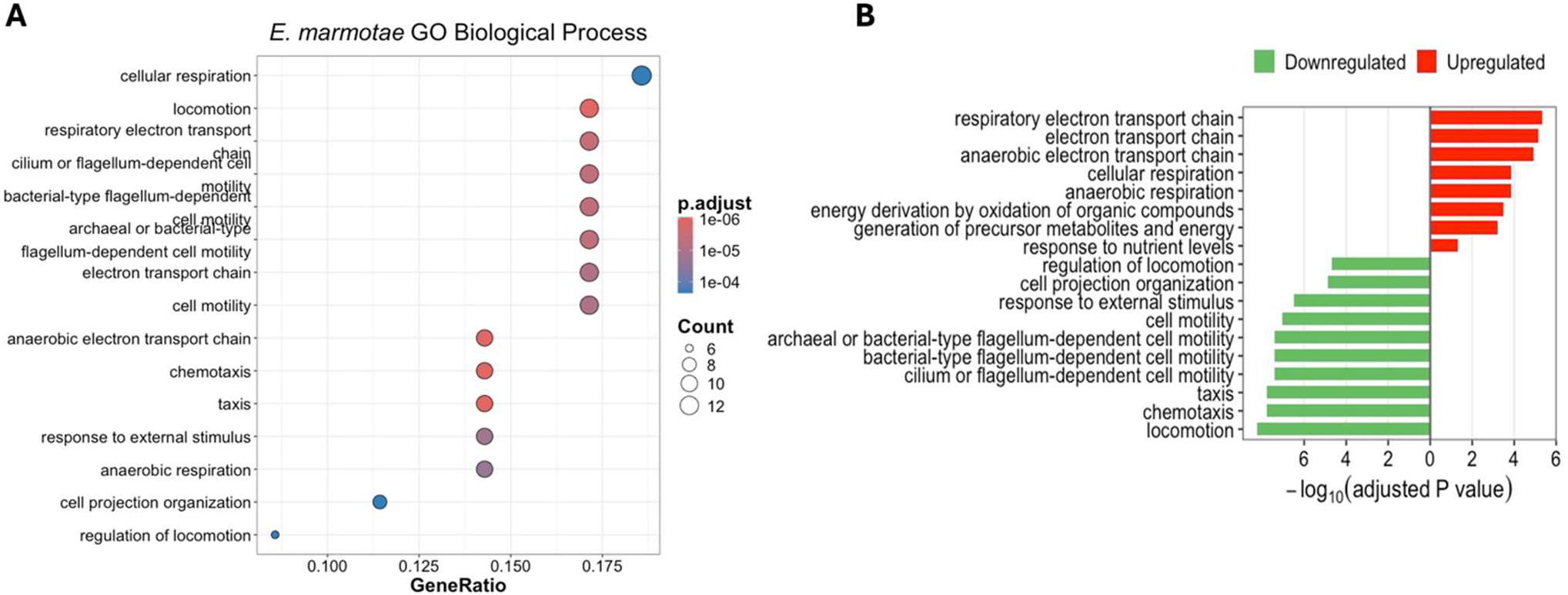
Gene Ontology (GO) enrichment analysis of differentially expressed genes in E. marmotae at 37 °C relative to 28 °C, showing significantly enriched categories of genes classified by Biological Process. (A) Dot size represents the number of genes associated with each term, while color indicates the adjusted p-values, according to the included scale. (B) Bars represent the adjusted p-values for downregulated gene categories (green), and upregulated gene categories (red). Bar length represents the enrichment significance expressed as -log_10_(adjusted p-value).

While all groups highlighted in Figure 6 by Gene Ontology (GO) biological process analysis for *E. marmotae* are significant at p < 0.05, we emphasize here those that are color-coded with the red and reddish-purple symbols (p < E-05) in Figure 6A, and the corresponding bars in Figure 6B. Processes with these highly significant adjusted p-values in *E. marmotae* include functional categories of locomotion, anaerobic electron transport, chemotaxis, and other taxis, with only slightly less significant p-values for several categories of flagellar-dependent motility. Of these process categories, all of those related to motility were significantly downregulated (Figure 6B) while the most significant upregulated categories were those related to electron transport and cellular respiration.

Comparable GO enrichment analysis in *E. coli* yields a completely different set of significantly enriched GO Biological Process differentially expressed categories. Figure 7 shows that differentially expressed gene categories were mainly associated with various amino acid metabolic processes (Figure 7A), and that these include many upregulated amino acid biosynthetic processes, while the significant downregulated categories include metabolic processes for organic acids, nitrates, ammonium, and other nitrogen cycle processes (Figure 7B).

**Figure 7.**
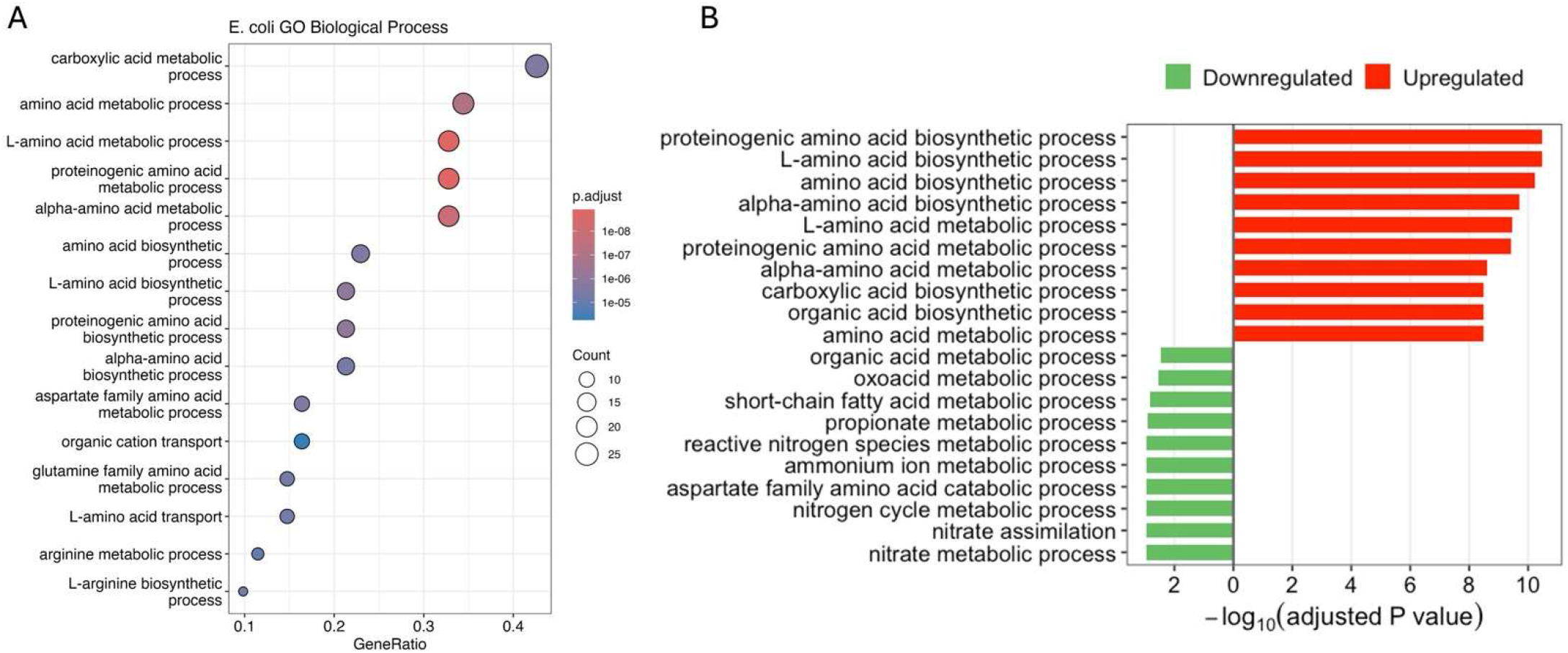
Gene Ontology (GO) enrichment analysis of differentially expressed genes in *E. coli* at 37 °C relative to 28 °C, showing significantly enriched categories of genes classified by Biological Process. (A) Dot size represents the number of genes associated with each term, while color indicates the adjusted p-values, according to the included scale. (B) Bar plot of the top enriched GO biological process terms among downregulated (green) and upregulated (red) gene categories. Bar length represents the enrichment significance expressed as -log_10_(adjusted p-value).

An informative alternate representation of functional groupings of differentially-expressed genes is using STRING network analysis, which takes into account not only the functions of their gene products used in ontology analysis, but also integrates information from a range of organisms on physical interactions of their gene products, systematic co-expression data, shared relationships across different organismal genomes, and relationship information mined from scientific literature. The results of STRING analysis of *E. marmotae*, illustrated in Figure 8, show that downregulated genes include an enormous “network ball” comprised of 21 genes involved with motility and their relationship partners. No network even closely resembling this was found in *E. coli* downregulated genes (Figure 9). Upregulated genes in the STRING analysis of *E. marmotae* differentially expressed genes are mostly in small isolated groups; however, one network with eight members is a cluster of *cdb* genes (cytochrome oxidase subunits; see https://string-db.org/network/511145.b0979) and *hya* genes (hydrogenase subunits, associated with cytochromes; see https://string-db.org/network/511145.b0975). This significantly upregulated cluster does not occur in *E. coli*; the major upregulated network (13 genes) in *E. coli* consists of many genes associated with arginine biosynthetic processes (*arg* genes and their associated network neighbors).

**Figure 8:**
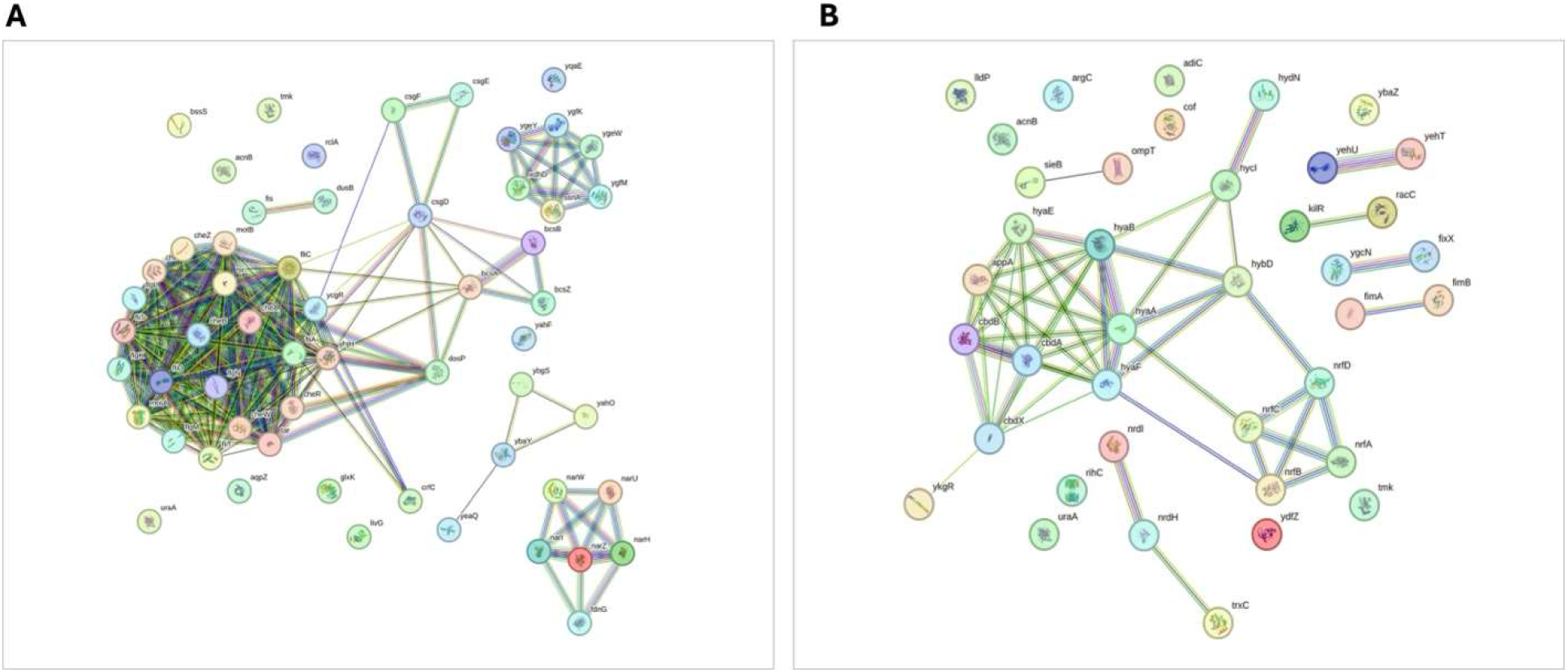
STRING network analysis of genes differentially expressed at 37 °C compared to 28 °C in *E. marmotae*. Strings connecting gene symbols represent known protein–protein interaction relationships of the gene products of (A) significantly downregulated genes and (B) significantly upregulated genes.

**Figure 9:**
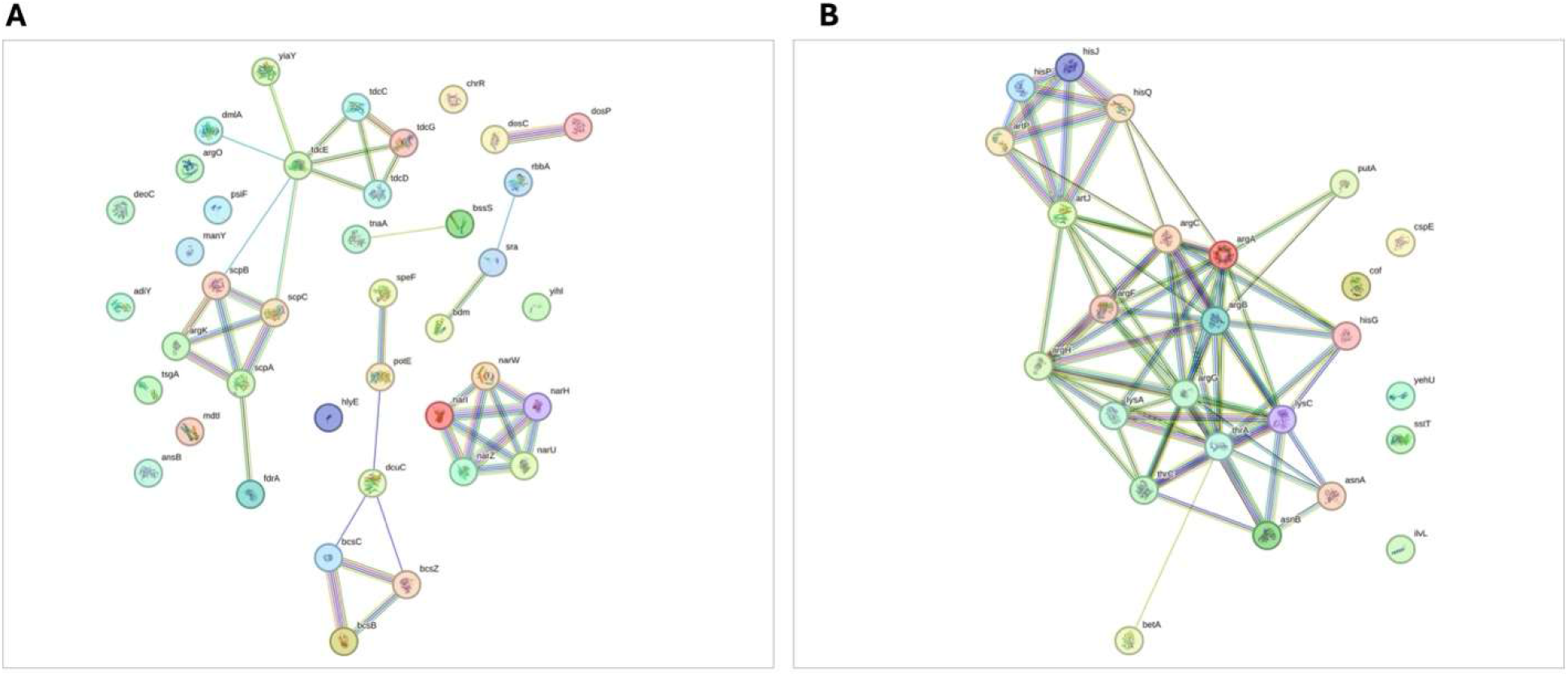
STRING network analysis of genes differentially expressed at 37 °C compared to 28 °C in *E. coli*. Strings connecting gene symbols represent known protein–protein interaction relationships of the gene products of (A) significantly downregulated genes and (B) significantly upregulated genes.

STRING analysis also highlighted a small network of temperature-responsive downregulated genes that was conserved between *E. marmotae* and *E. coli.* In Figure 8A and Figure 9A, both species exhibited an isolated “pentagon-shaped network” with an embedded start-shaped string pattern at the lower right of each of these downregulation illustrations. These networks consist of gene products that are coordinately downregulated to reduce expression of genes involved in nitrate reduction (*narZ/U/H/W/I*) and an additional gene in *E. marmotae, fdnG*, a nitrate-inducible formate dehydrogenase known to function in electron transport when nitrate is present.

Overall, these results indicate that in *E. marmotae*, higher temperature is associated with a pronounced repression of motility genes, while *E. coli* exhibited stronger enrichment of metabolic adjustment pathways. These observations are consistent with previous studies demonstrating temperature-dependent regulation of motility-associated genes in *E. marmotae* (15).

### 3.2. Differential expression of proteins

Proteomic analyses were conducted on *E. marmotae* grown under identical conditions as above to assess the effect of temperature on protein amounts. Consistent with the transcriptomic data, an overall comparison revealed clear thermoregulatory trends in the proteome. Of the 2,900 proteins detected (Table S3), 110 proteins were differentially present in response to temperature, using a cutoff of ±1 Log₂ fold change (Figure 10 and Table S3). Among these differentially affected proteins, the majority (80 proteins) were decreased at 37 °C relative to 28 °C.

**Figure 10:**
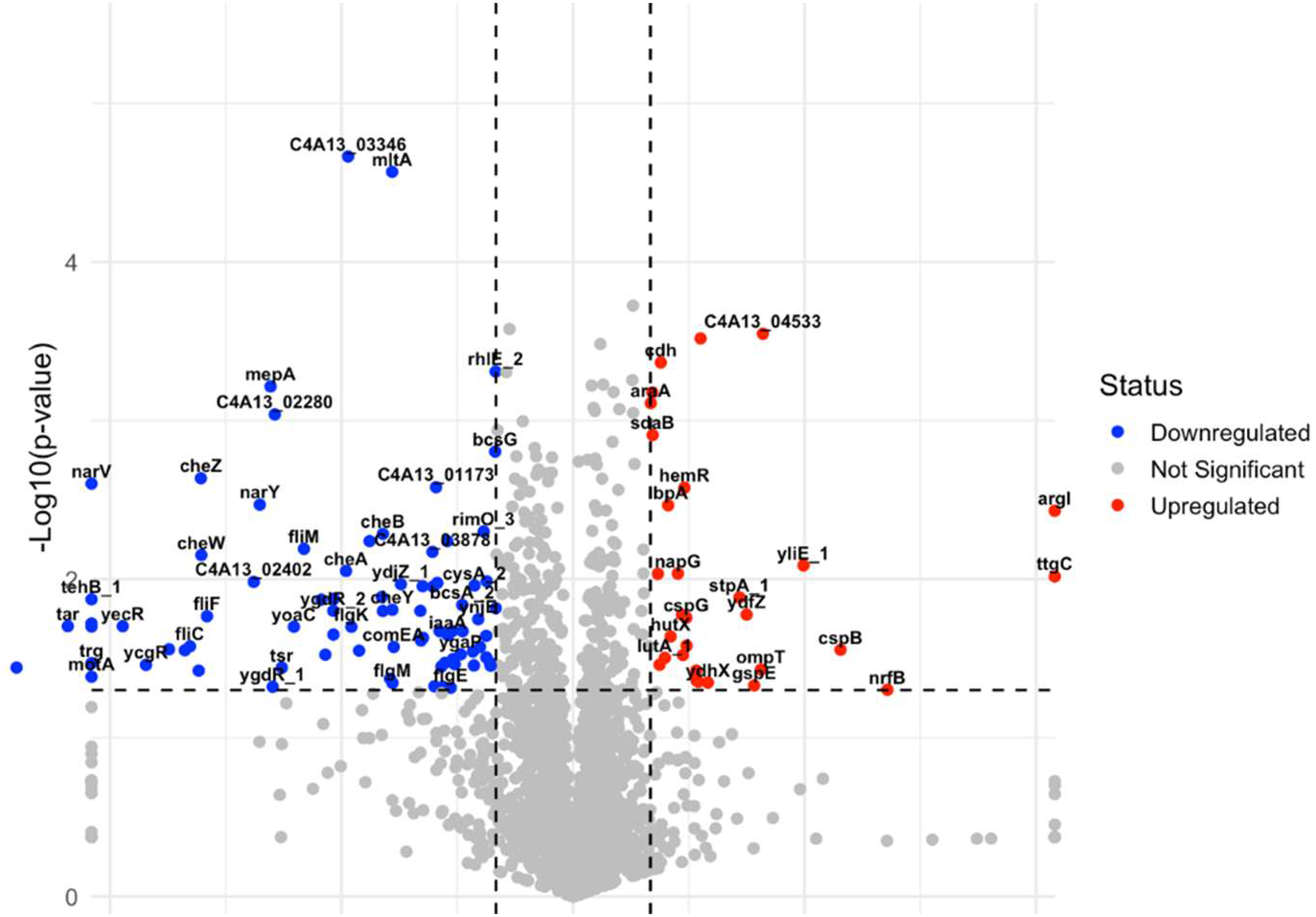
Volcano plot showing differential protein amounts in *E. marmotae* at 37 °C versus 28 °C. Proteins significantly reduced (blue) or increased (red) in amount are shown based on a ±1 Log₂ fold change and –log₁₀(p-value) threshold. Numerous motility- and chemotaxis-associated proteins are significantly downregulated at elevated temperature.

A substantial proportion of the decreased proteins are associated with bacterial motility and chemotaxis. Several key motility-related proteins, including motility protein A (*motA*), flagellar biosynthetic protein *fliP*, flagellar protein *fliO*, the aerotaxis receptor *Aer*, methyl-accepting chemotaxis protein III (*trg*), and *Yjcz*, were detected at 28 °C but were not detected at 37 °C. Comparison of mRNA expression and protein amount levels revealed strong agreement for many genes, particularly those involved in motility and chemotaxis, which were consistently reduced at both the transcript and protein levels (Figure 11).

**Figure 11:**
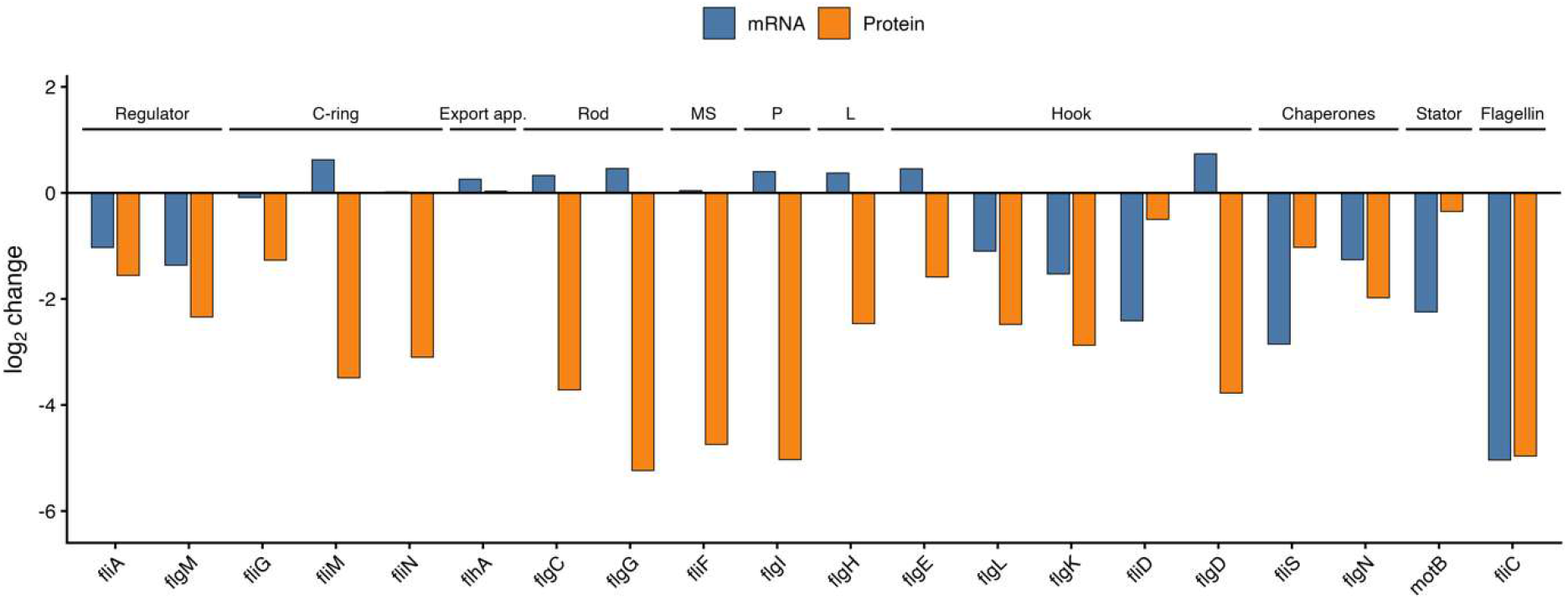
Comparison of log₂ fold changes in mRNA (blue bars) and protein abundance (orange bars) for genes associated with flagellar assembly, motility, chemotaxis, chaperones, and stress response, for *E. marmotae* when grown at 37 °C relative to 28 °C.

However, notable discrepancies between transcriptomic and proteomic data were also observed. Several components of the flagellar ATPase complex, rod, hook, and P- and L-rings exhibited markedly stronger decreases in the amount of protein compared to the decreases in the mRNA level (Figure 11). In some cases, these genes were not significantly downregulated at the transcript level, despite pronounced reduction of the corresponding proteins. These results may indicate that post-transcriptional regulation and/or protein stability play an important role in the temperature-dependent control of flagellar assembly in *E. marmotae*.

In contrast, proteins that increased at 37 °C were predominantly associated with stress adaptation, anaerobic respiration, nutrient acquisition, and host interaction. Given that 37 °C serves as an environmental cue for entry into the human host, many bacterial virulence-associated genes are known to be thermoregulated (18, 31). Among the proteins increased at 37 °C was OmpT, an outer membrane protease that has been identified as a virulence factor in most uropathogenic *E. coli* (UPEC). *ompT* contributes to UPEC pathogenesis by promoting adhesion to human epithelial cells and by cleaving host antimicrobial peptides, thereby facilitating bacterial colonization, invasion, and immune evasion (32, 33). Deletion of *ompT* in enteropathogenic and uropathogenic *E. coli* results in reduced colonization and virulence (33). The increased expression of *ompT* at both the transcriptomic and proteomic levels in *E. marmotae* at 37 °C suggests that *E. marmotae* enhance *ompT* production in response to host-associated temperature, potentially contributing to adaptation and virulence during infection. Although the role of *ompT* in *E. marmotae* pathogenesis remains unknown, its thermoregulated expression, together with its established function in pathogenic *E. coli*, suggests that it may contribute to the ability of *E. marmotae* to establish infections such as urinary tract infections and sepsis. These findings warrant further functional studies to determine the contribution of *ompT* to *E. marmotae* virulence.

The above results were compared to *E. coli* samples grown under identical conditions to assess the effect of temperature on protein amounts. Consistent with the transcriptomic data, an overall comparison revealed clear thermoregulatory trends in the proteome. Of the 2,980 proteins detected (Table S4), 45 proteins were differentially changed in response to temperature, using a cutoff of ±1 Log₂ fold change (Figure 12; Table S4). Among these differentially present proteins, the majority of 22 proteins were downregulated at 37 °C relative to 28 °C. The proteomics analysis showed coordinated upregulation of the nitrate and nitrite reductase-associated proteins NarG/H/I and NrfAC.

**Figure 12:**
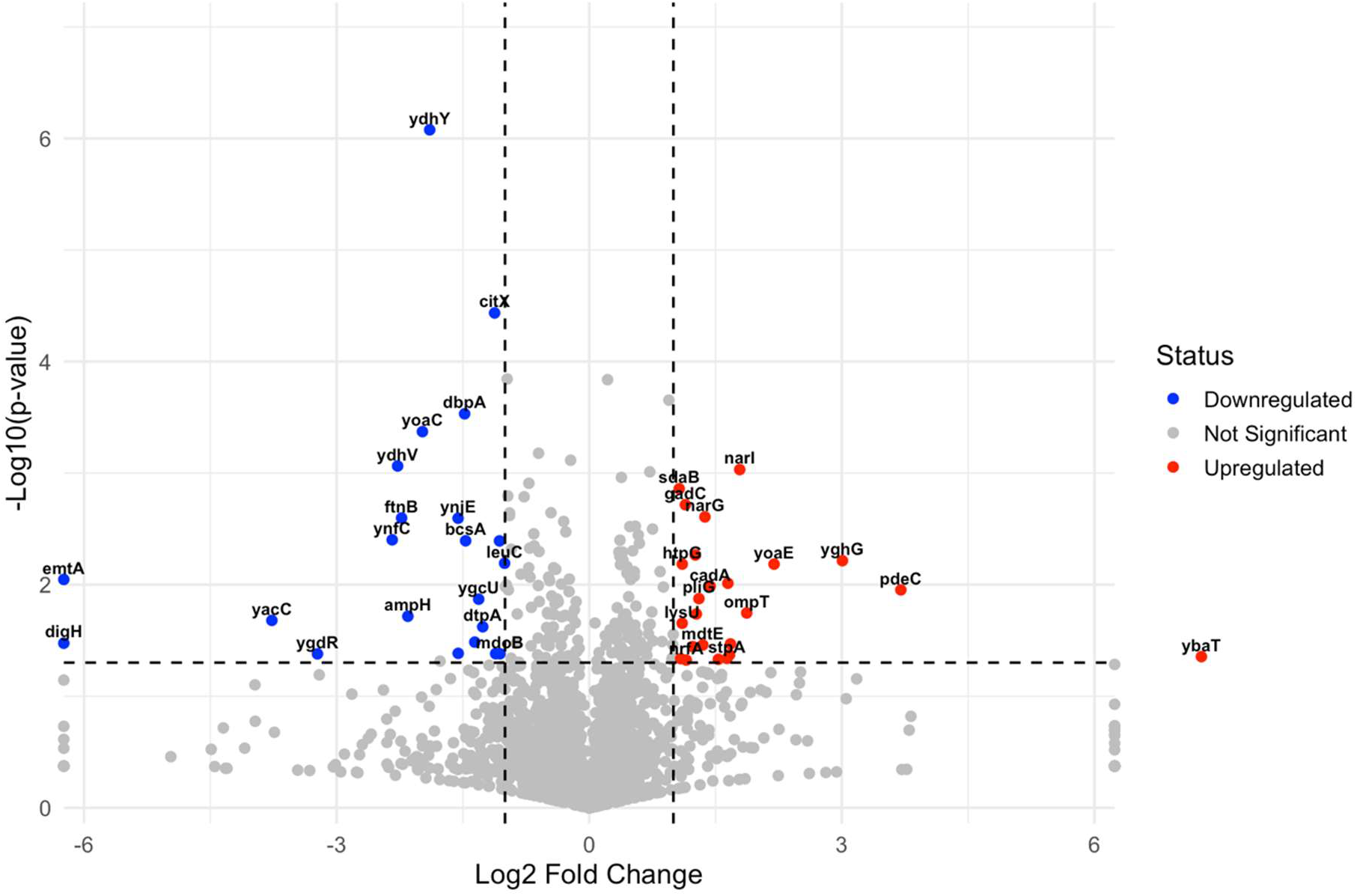
Volcano plot showing differential protein expression of *E. coli* at 37 °C versus 28 °C. Proteins significantly downregulated (blue) or upregulated (red) are shown based on a ±1 Log₂ fold change and –log₁₀(p-value) threshold. Numerous motility- and chemotaxis-associated proteins are significantly downregulated at elevated temperature.

A second important group includes genes involved in acid resistance and stress adaptation, such as *gadC, cadA, htpG, lysU, aidB, speG,* and *yqhD*. The upregulation of GadC and CadA protein levels is especially relevant because glutamate- and lysine-dependent acid resistance systems help enteric bacteria survive acidic environments. Several increased proteins also point toward host interaction and survival under antimicrobial pressure. The *ompT* gene encodes an outer membrane protease associated with cleavage of antimicrobial peptides and immune evasion (34, 35), while *mdtE* encodes part of a multidrug efflux system that can contribute to resistance against toxic compounds (36, 37). These changes may indicate that 37 °C acts as a host-temperature signal that enhances protective mechanisms relevant to infection.

## 4.0 Discussion

Overall, our findings suggest that flagellar motility is the major cellular function that is downregulated by *E. marmotae* at human body temperature. The transcriptional and phenotypic data together support a model in which motility in *E. marmotae* is strongly repressed at 37 °C relative to 28 °C. This interpretation is consistent with the observed reduced motility phenotype at 37 °C and with the coordinated downregulation of genes involved in flagellar assembly, motor function, and chemotaxis, including *fliA*, *fliC*, *motA*, *motB*, *cheA*, and *cheY* (15). Together, these data suggest that the loss of motility at 37 °C is not due to isolated gene-level changes, but rather reflects broad repression of the flagellar and chemotaxis regulatory programs.

The downregulation of *fliA* is particularly important because *fliA* encodes the flagellar sigma factor σ^28^, which activates late-stage flagellar genes required for filament formation, motor activity, and chemotaxis. Reduced *fliA* expression therefore provides a plausible regulatory explanation for the decreased expression of downstream structural and chemotaxis components (38, 39). In contrast, expression of the genes encoding the master flagellar regulator, *flhDC,* were not significantly altered, suggesting that temperature-dependent motility repression in *E. marmotae* may occur primarily at the level of the late flagellar regulon rather than through complete shutdown of the uppermost flagellar regulatory hierarchy. This distinction is important because it indicates a targeted remodeling of motility gene expression at 37 °C.

Among the top 20 significantly downregulated genes in *E. marmotae* is *pdeH*, whose gene product is an EAL domain phosphodiesterase that may provide a second regulatory layer for motility. Its phosphodiesterase activity lowers intracellular cyclic-di-GMP, a conserved bacterial second messenger that controls the transition between motile and sessile lifestyles. In many bacteria, low intracellular cyclic-di-GMP generally favors motility, whereas elevated cyclic-di-GMP promotes surface attachment, biofilm formation, and reduced motility (40, 41). Therefore, the downregulation of *pdeH* (also known as *yhjH*) (42)) in *E. marmotae* at 37 °C supports a model in which altered cyclic-di-GMP homeostasis reinforces the non-motile phenotype. This is consistent with previous studies showing that deletion of *yhjH* increases c-di-GMP levels and suppresses motility, while deletion of the c-di-GMP effector *ycgR* can partly restore motility in a *yhjH* mutant background (43, 44). Therefore, we postulate that in *E. marmotae*, reduction of *pdeH* transcription and its resultant gene product activity can increase cyclic-di-GMP levels and shift cells toward a less motile condition.

However, the expression pattern of *ycgR* suggests that the 37 °C phenotype is unlikely to be driven primarily by a *ycgR*-mediated motor-braking mechanism. YcgR is a PilZ-domain cyclic-di-GMP effector that can act as a flagellar brake in *E. coli* and *Salmonella* by interacting with motor-associated proteins such as FliG, FliM, and MotA when cyclic-di-GMP levels are elevated (45). In the present dataset, however, *ycgR* was also downregulated at 37 °C. This argues against a model in which repression of motility is mainly caused by an active YcgR-dependent brake acting on an otherwise assembled and functional flagellar motor. Instead, the broad downregulation of *fliA*, *fliC*, *motA*, *motB*, *cheA*, and *cheY* indicates that the motility apparatus itself is transcriptionally curtailed. Thus, reduced *pdeH* expression remains compatible with a shift toward higher cyclic-di-GMP and reduced motility, but the dominant mechanism appears to be transcriptional repression of the late flagellar and chemotaxis programs, with cyclic-di-GMP signaling acting as a reinforcing rather than primary mechanism.

The accompanying GO enrichment results strongly reinforce this interpretation. Downregulated genes in *E. marmotae* were enriched in processes such as bacterial-type flagellum-dependent cell motility, locomotion, cell projection organization, and response to external stimulus. These enriched categories closely mirror the gene-level differential expression data and confirm that the 37 °C response is not driven by isolated genes, but by coordinated suppression of an entire functional network. By contrast, upregulated genes were enriched in processes related to cellular respiration, response to nutrient levels, and anaerobic electron transport, suggesting that elevated temperature may also promote physiological adjustment in energy generation and nutrient utilization. Thus, the *E. marmotae* transcriptome at 37 °C appears to shift away from active environmental exploration and toward a state less dependent on motility and more focused on metabolic adaptation.

Indeed, genes upregulated at 37 °C were enriched in processes related to cellular respiration, response to nutrient levels, and anaerobic electron transport. Elevated temperature may promote broader physiological remodeling, including changes in energy generation, nutrient utilization, and adaptation to host-associated conditions. Therefore, the transcriptome of *E. marmotae* at 37 °C appears to reflect a transition from an environmentally exploratory state toward a less motile and more host-adapted physiological state. Such a shift may be advantageous during infection, where reduced motility can help bacteria conserve energy, avoid immune recognition associated with flagellin, and prioritize traits involved in persistence, immune evasion, and survival in host tissues.

In addition to repression of motility, several genes with potential roles in host interaction and immune evasion showed increased expression at 37 °C. These included genes associated with outer membrane protease *ompT*, encoding type 1 fimbrial proteins (*fimA*, and *fimB* ) . Among these upregulated genes, increased levels of their protein products in *E. marmotae* included OmpT. The outer membrane protease OmpT also emerged as one of the strongly thermoregulated molecules in both the transcriptome and proteome (Figures 2 and 10). OmpT is a recognized virulence-associated protease in pathogenic *E. coli* and contributes to immune evasion by cleaving antimicrobial peptides and proteins, including cathelicidin (LL-37) (35), protamine (46), bacterial colicins E2, E3, and D (47), and antimicrobial proteins isolated from human urine (48). OmpT has also been implicated in adhesion, invasion, intracellular bacterial community formation, and induction of host inflammatory cytokines in uropathogenic *E. coli* (32). Its increased expression at 37 °C in *E. marmotae* therefore suggests that this organism may enhance specific immune evasion and host-interaction mechanisms under human body temperature conditions. Together, these findings suggest that *E. marmotae* may reduce energetically costly motility while increasing factors that promote persistence, immune evasion, and host adaptation.

The expression of fimbrial genes provides another important dimension to the temperature-dependent response. Type 1 fimbriae are major adhesins in bacteria and are central to host attachment, biofilm development, and persistence during infection (49, 50). *fimA* encodes the major structural subunit of the type 1 pilus rod and supports pilus assembly, which enables bacterial attachment to mannose-containing receptors on epithelial surfaces, including the uroepithelium (51, 52). In this study, the differential expression of *fimA/fimB* suggests that *E. marmotae* may regulate adhesive structures in response to host temperature. This is biologically relevant because repression of motility and increased expression of adhesive or surface-associated factors often occur together during bacterial transition from environmental dispersal to host colonization. However, *fimH*, which encodes the mannose-binding adhesin at the tip of type 1 fimbriae and is a key determinant of uropathogenic *E. coli* attachment during urinary tract infection, was not significantly differentially expressed.

The increased expression of *cdtB* (Table S2) is also notable because cytolethal distending toxin B is the active DNase-like subunit of the CDT toxin complex. CDT-producing bacteria can induce DNA damage, cell cycle arrest, and apoptosis in eukaryotic cells, with *cdtB* mediating much of this genotoxic activity (53). The presence and increased expression of *cdtB* at 37 °C may therefore indicate a potential mechanism by which *E. marmotae* could damage host cells or modulate host tissue responses during infection. Although this finding is intriguing, it should be interpreted cautiously until toxin production, secretion, and host-cell effects can be experimentally investigated.

Relatively few studies have examined the effect of temperature on global gene expression in *E. coli*, and those that do have generally compared temperatures different from those used in the present study (54–56). Gant Kanegusuku et al. (2021) compared gene expression at 23 °C and 37 °C, whereas Rudenko et al. (2019) examined responses to heat stress by comparing 37°C and 42°C. These larger temperature shifts are expected to induce broader physiological responses than the environmentally relevant transition from 28 °C to 37 °C investigated here.

Consistent with previous studies, *ompT* expression increased in *E. coli* at 37 °C, suggesting that thermoregulation of this virulence-associated protease is conserved among pathogenic and emerging pathogenic *Escherichia* species (54, 55). However, several important differences were also observed. Gant Kanegusuku et al. (2021) reported reduced curli expression and biofilm formation at 37 °C relative to 23 °C, whereas our study identified significant downregulation of cellulose biosynthesis genes (*bcsB, bcsC,* and *bcsZ*) and little evidence for major changes in curli-associated pathways between 28 °C and 37 °C. Likewise, previous studies reported increased expression of flagellar and motility-associated genes at 37 °C relative to 23 °C, whereas no significant temperature-dependent regulation of motility was observed in *E. coli* under our experimental conditions.

Taken together, these results support a temperature-dependent adaptive model for *E. marmotae*. At 28 °C, a temperature more consistent with environmental reservoirs, *E. marmotae* maintains higher expression of motility and chemotaxis genes, supporting movement, dispersal, and environmental exploration. At 37 °C, a temperature associated with mammalian host infection, the organism strongly represses the late flagellar and chemotaxis regulon, resulting in a non-motile phenotype that may provide an advantage during host association by reducing immune recognition of flagellin and limiting activation of flagellin-mediated innate immune responses. This shift away from motility may also allow the organism to conserve energy and redirect cellular resources toward traits that promote persistence, stress tolerance, and survival under host-like conditions.

On the other hand, the loss of motility may make some infection sites and routes less accessible to *E. marmotae* than to *E. coli*, for example. Retrograde motility-driven invasion of the kidneys (49, 57) may be less likely; however, uropathogenic infections in which flagella are suppressed to avoid immune responses may be an even more important mechanism in maintaining urinary tract infections. Further studies, with recently developed diagnostic tools (9) may help determine if the spectrum of infections and infection etiology differs between *E. marmotae* and *E. coli*.

Overall, our findings suggest that temperature is a key regulator of *E. marmotae* biology and may help explain how this organism transitions from environmental reservoirs to human infection. The coordinated repression of motility genes, together with the increased expression of selected host-associated and virulence-related factors, supports a model in which *E. marmotae* adapts to mammalian temperature by reducing traits associated with environmental dispersal while enhancing features that may support immune evasion and pathogenic potential. This temperature-dependent regulatory strategy provides important insight into the emergence of *E. marmotae* as an underrecognized human pathogen and highlights the need for future studies to directly test the roles of cyclic-di-GMP signaling, outer membrane proteins, fimbrial structures, and toxin-associated genes in host colonization and virulence.

## DECLARATIONS

### Availability of data and materials

The raw RNA sequencing data have been deposited in the NCBI Sequence Read Archive (SRA) under accession number SUB16381930. The mass spectrometry proteomics data have been deposited in the ProteomeXchange Consortium via the PRIDE repository under accession PXD081413. The whole-genome sequence assemblies of the *Escherichia marmotae* RAM laboratory isolates used in this study were previously published (Oladipo et al., 2025a) and are available in GenBank under accession numbers **J**BNVMU000000000 (RAM 3032), JBNVMW000000000 (RAM 3054), and JBNVMX000000000 (RAM 3024).

### Consent for publication

All authors contributed to writing – review & editing and approved the final version of the manuscript.

## Abbreviations

LB: Luria-Bertani
WGS: Whole genome sequencing
MALDI-TOF MS: Matrix-Assisted Laser Desorption Ionization-Time of Flight Mass Spectrometry
UTI: Urinary tract infection
PBS: Phosphate-Buffered Saline
ABC: ATP-binding cassette transporter
GO: Gene Ontology
FDR: False Discovery Rate
PCA: Principal Component Analysis
c-di-GMP: cyclic di-guanylate (bis-(3′-5′)-cyclic diguanylic acid)
PPIN: Protein-protein Interaction Network
UPEC: Uropathogenic *<u>E.coli</u>*
GSEA: Gene Set Enrichment Analysis
COG: Clusters of Orthologous Groups
IDT: Integrated DNA Technologies

